# High-Resolution Spatial Profiling of Microglia Reveals Proximity Associated Immunometabolic Reprogramming in Alzheimer’s Disease

**DOI:** 10.1101/2025.05.16.654329

**Authors:** Kai Saito, Danielle S. Goulding, Georgia L. Nolt, Sophia H. Dimas, Lauren C. Moore, Isaiah O. Stevens, Sonya Anderson, Andy Snipes, Shannon L. Macauley, Peter T. Nelson, Lance A. Johnson, Josh M. Morganti

## Abstract

Single-cell RNA sequencing has demonstrated that the presence of parenchymal amyloid plaques and intracellular hyperphosphorylated tau pathology is associated with distinctive (and possibly disease-driving) microglial heterogeneity. However, our understanding of how proximity to these Alzheimer’s disease (AD) pathological hallmarks in situ relates to microglial gene expression remains obscure. Here, we utilized high-resolution spatial transcriptomics (ST) via the Xenium platform with a fully customized gene panel to elucidate disease-associated microglial subtypes in tandem with examining metabolic signatures across AD-relevant mouse models and well-characterized human postmortem tissue. Three mouse models were evaluated: PS19, APP/PS1, and 5xFAD. Analyzing anatomical features across entire hemisections, our approach resolved the distribution of five disease-associated microglial subtypes, while deciphering how proximity to cerebral amyloid plaques influenced transcriptional mediators governing metabolic pathways. We observed robust alterations in glycolytic and cholesterol/lipid processing pathways in plaque-associated microglia, consistent with a specific switch to glycolysis and lipid-fueled metabolism in the plaque niche. Extending our analysis to human postmortem dorsolateral prefrontal cortex (dlPFC), we identified conserved disease-reactive microglial states, i.e., similar proximity-dependent metabolic shifts around amyloid plaques. Further, integrating spatial transcriptomics with machine-learning approaches revealed novel anatomic domain-specific cellular gene expression profiling features, highlighting differential vulnerabilities of neuronal, glial, and vascular populations near specific microglial subtypes. Together, our findings provide one of the first comprehensive and high-resolution atlas of microglial immunometabolic states across species, anatomical regions, and AD pathological burden.

## Introduction

Microglia constitute the brain’s largest population of resident immune cells serving a variety of dynamic roles throughout the lifespan. In the context of aging and diseased brains, a multitude of microglial phenotypes have been described with increasing clarity due to expanding capabilities harnessing single cell resolving proteomic and transcriptomic approaches ^1-9^. Particular attention has been spent examining microglial reactive states in various animal models recapitulating pathophysiological elements of AD and AD-related dementia (ADRD). Pathoetiological susceptibility to and progression of AD/ADRD have been linked to microglial responses via multiple genome-wide association studies uncovering AD/ADRD risk loci being preferentially expressed by microglia. Interestingly, many of these genes have direct links to cellular metabolism^10-13^, suggesting mechanism(s) in disease-associated immunometabolic reprogramming of microglia. Consistent with these insights from genetic studies, single-cell transcriptomic studies have identified distinct disease-associated microglial subpopulations in both AD-like animal models as well as post-mortem AD brain tissue that exhibit hallmarks of immunometabolic reprogramming ^4,5,14,15^. Together, these transcriptomic findings bridge AD risk gene activity with microglial functional states, underscoring immunometabolic remodeling as a hallmark of the disease-associated microglial phenotype. However, despite advances in defining microglial states, our understanding of spatial context in microglial heterogeneity remains incomplete. Most single-cell transcriptomic and cytometric studies rely on dissociating cells from their tissue environment, which inherently loses information regarding the impact of how microenvironments, anatomy, and proximity to pathology may shape microglial phenotypes.

Until recently, we lacked the tools to profile microglial gene expression in situ with high resolution. Traditional histological or imaging approaches could map a few markers in tissue, but they did not capture the broader transcriptomic programs of microglia nor quantitatively link those programs to distance from pathological landmarks. This gap is beginning to be addressed by integrative spatial ‘omics technologies^16,17^. High-resolution or single cell resolving spatial transcriptomics now allows transcript profiling directly on tissue sections, preserving spatial coordinates for each measured cell. Recent work has begun to demonstrate that cellular cross-talk niches exist between microglia and astrocytes, which is altered by the presence of amyloid burden ^18^. Still, spatial transcriptomic analyses of microglia are only in their infancy, and important knowledge gaps remain – including how metabolic reprogramming manifests across different brain regions and disease stages, and whether spatially-defined microglial states observed in mice correspond to similar states in human AD brains. To date, spatial profiling studies have been limited by optical resolution requiring deconvolution to impute cell-type abundance, and narrow area of coverage often encompassing a single model or discrete brain region, leaving many questions about microglial state diversity in situ unanswered.

Here, we sought to overcome these limitations by leveraging high-resolution spatial transcriptomics to map microglial immunometabolic states directly within intact brain sections from multiple AD models and human patients. We used the 10x Genomics *Xenium* in situ platform with a customized gene panel enriched for immune and metabolic genes, enabling the multiplex detection of transcripts at single-cell (and subcellular) resolution in tissue. Using this approach, we profiled microglial gene expression landscapes in three complementary AD mouse models – 5xFAD and APP/PS1 mice (which develop amyloid plaques) and PS19 tau transgenic mice (which develop tau neurofibrillary pathology) – as well as in postmortem dorsolateral prefrontal cortex (dlPFC) from Braak VI AD tissue. This strategy allowed us to examine microglial phenotypic diversity across different types of AD-related lesions (amyloid vs. tau), across species, and within spatially defined neighborhoods relative to AD-related pathological burden.

Our spatially resolved transcriptomic atlas reveals that microglia in AD brains segregate into multiple transcriptional subtypes with disease-associated profiles. We found that disease-associated microglial subtypes were enriched in proximity to amyloid plaques forming niches of reactive microglia in the tissue. Microglia occupying plaque-rich regions showed markedly elevated expression of glycolytic enzymes and lipid metabolism genes, consistent with a switch to glycolysis and lipid-fueled metabolism in the plaque niche. By contrast, microglia located farther from plaques tended to retain homeostatic and/or less metabolically active profiles. Irrespective of mathematically-defined states/clusters, spatial proximity to pathology emerged as a strong predictor of microglial transcriptional state: the nearer a microglial cell was to a plaque, the more it upregulated genes for glucose utilization, lipid catabolism, and pro-inflammatory mediators. This spatial pattern of microglial metabolic reprogramming was conserved across species – we observed strikingly similar plaque-associated microglial states in both the mouse models and human AD cortex. Furthermore, integrative analysis of the tissue microenvironment identified distinct cellular “neighborhoods” in which activated microglia co-localize with reactive astrocytes and other glial cells around pathology, highlighting specific niches of neuroinflammatory metabolism. Taken together, our study provides a comprehensive *in situ* view of microglial heterogeneity in AD, demonstrating how anatomical context and disease milieu shape microglial immunometabolic states. These findings fill an important gap between single-cell data and histopathology, and they lay the groundwork for spatially targeted therapeutic strategies aiming to modulate microglial metabolism in Alzheimer’s disease.

## Results

### Alzheimer’s disease related pathology drives spatial heterogeneity of microglia

In our mouse models, we acquired spatial transcriptional location of 797,255 cells across a 12 coronal sections, comprised of 19 clusters. Though our principal goal was to examine microglial-specific spatial repertoire, we were able to define multiple additional CNS cell types such as excitatory and inhibitory neurons, oligodendrocytes, oligodendrocyte precursor cells (OPCs), astrocytes, endothelial cells, pericytes, vascular leptomeningeal (VLMCs), lymphocytes, monocytes/macrophages, ependymal, and choroid plexus populations (Figure 1A-D). Cell typing across these canonical class levels revealed a rational spatial organization of relevant cell types across all sections. For example, white matter tracts comprising the corpus callosum were heavily enriched in oligodendrocyte-labeled cells and excitatory neurons comprising the CA1-3 and dentate gyrus of the hippocampus, VLMCs were observed forming boundaries, and ependymal cells clearly lining ventricular cavities were readily identifiable in a highly resolved manner across all sections (Figure 1D-F). Therefore, our panel, while not exhaustive of 100’s of genes to define various CNS resident cells was still able to reliably parse out relevant CNS cellular architecture.

**Figure 1.**
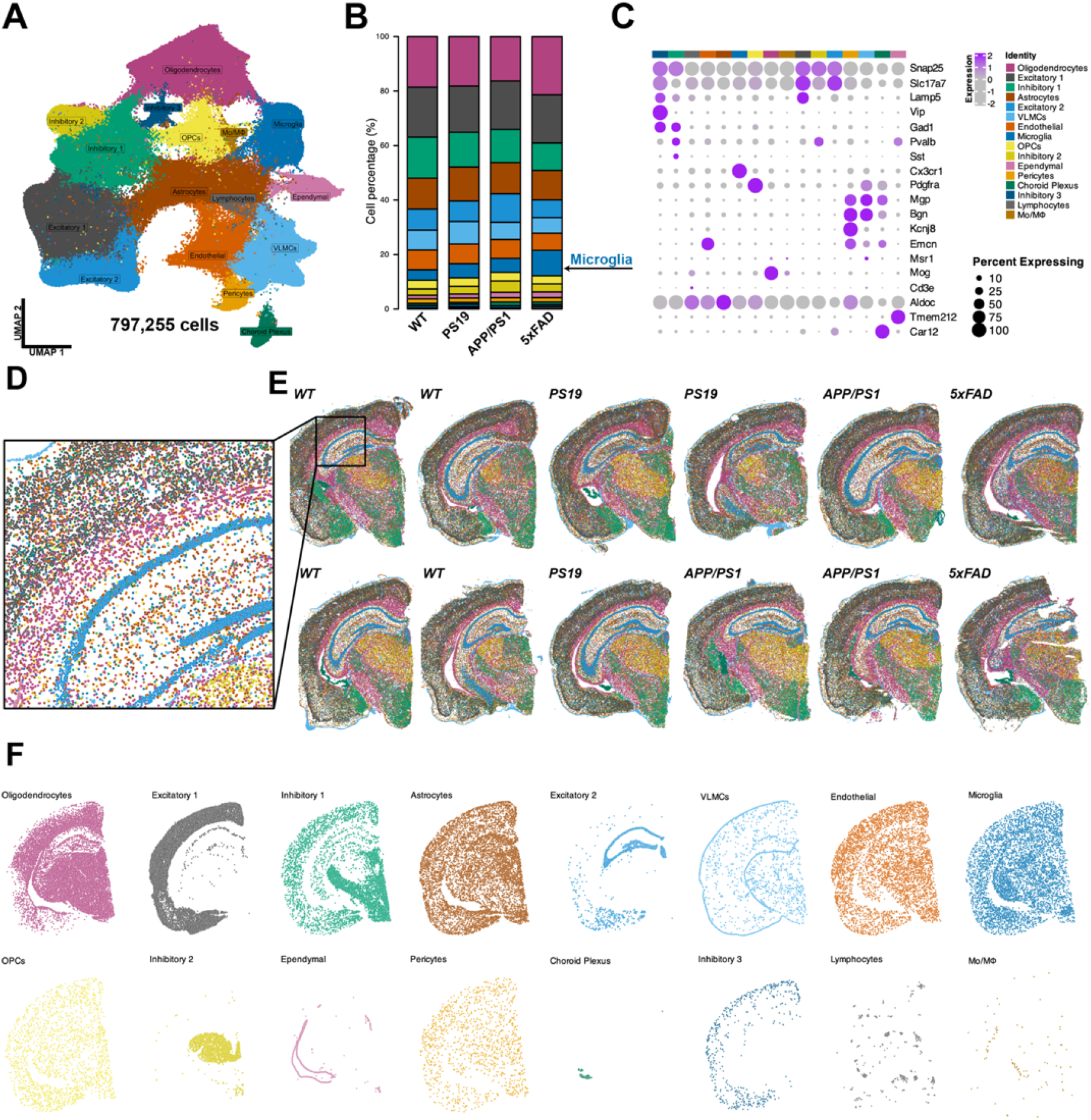
Single-cell resolving spatial transcriptomics reveals diverse landscape of sixteen principal cell classes. **A.)** UMAP of 797,255 classified cell types from aggregated samples (n=12 coronal sections) annotated by 16 cell classes. **B.)** Stacked bar plot on relative proportions across WT (n=4), PS19 (n=3), APP/PS1 (n=3) and 5xFAD (n = 2), with microglia highlighted in blue. **C.)** Scaled dot plot of canonical cell type marker genes used to define each cluster. **D.)** High-resolution crop of the dorsal hippocampus from a WT replicate demonstrating the single-cell resolving spatial location of multiple classes of cells, for example; excitatory (gray and blue), oligodendrocytes (maroon), inhibitory (yellow) are clearly delineated in anatomically rational locations. **E.)** Whole coronal genotype replicates for the twelve samples analyzed showing sixteen cell classes. **F.)** Representative 5xFAD coronal replicate split by sixteen cell classes to demonstrate anatomically and regionally enriched locales.

Next, we examined microglial responses beyond the broad class-level identifiers for cellular heterogeneity across our 4 genotypes. We quantified a total of 39,058 microglia between our 12 samples identifying 9 states: Mg1-9 (Figure 2A). Proportionally, Mg1-4 predominated in the WT mice, while Mg5-9 were proportionally expanded to varying levels across PS19, APP/PS1, and 5xFAD genotypes (Figure 2B). Comparatively, 5xFAD tissue acquired the principal changes across cell type distributions in our dataset, with significant decreases in Mg1 and Mg4 and concomitant increases in Mg5-9, relative to WT levels. Similarly, both APP/PS1 and PS19 had significant increases in Mg5-6 subtypes, with PS19 also having significantly more Mg8, compared to WT tissues. Grossly, the finding of dichotomy between non-diseased versus disease-reactive microglia states recapitulates many prior scRNAseq datasets ^4-6,14,19-21^. When we examined differentially expressed markers per each of the 9 states, we observed that the 4 states enriched in WT mice (i.e. Mg1-4) had the highest expression of ‘homeostatic’ associated markers *Tmem119, P2ry12, and Selpg*, contrasting with Mg5-9, which were comparatively downregulated as these were the states predominantly enriched across the three AD disease models (Figure 1C).

**Figure 2.**
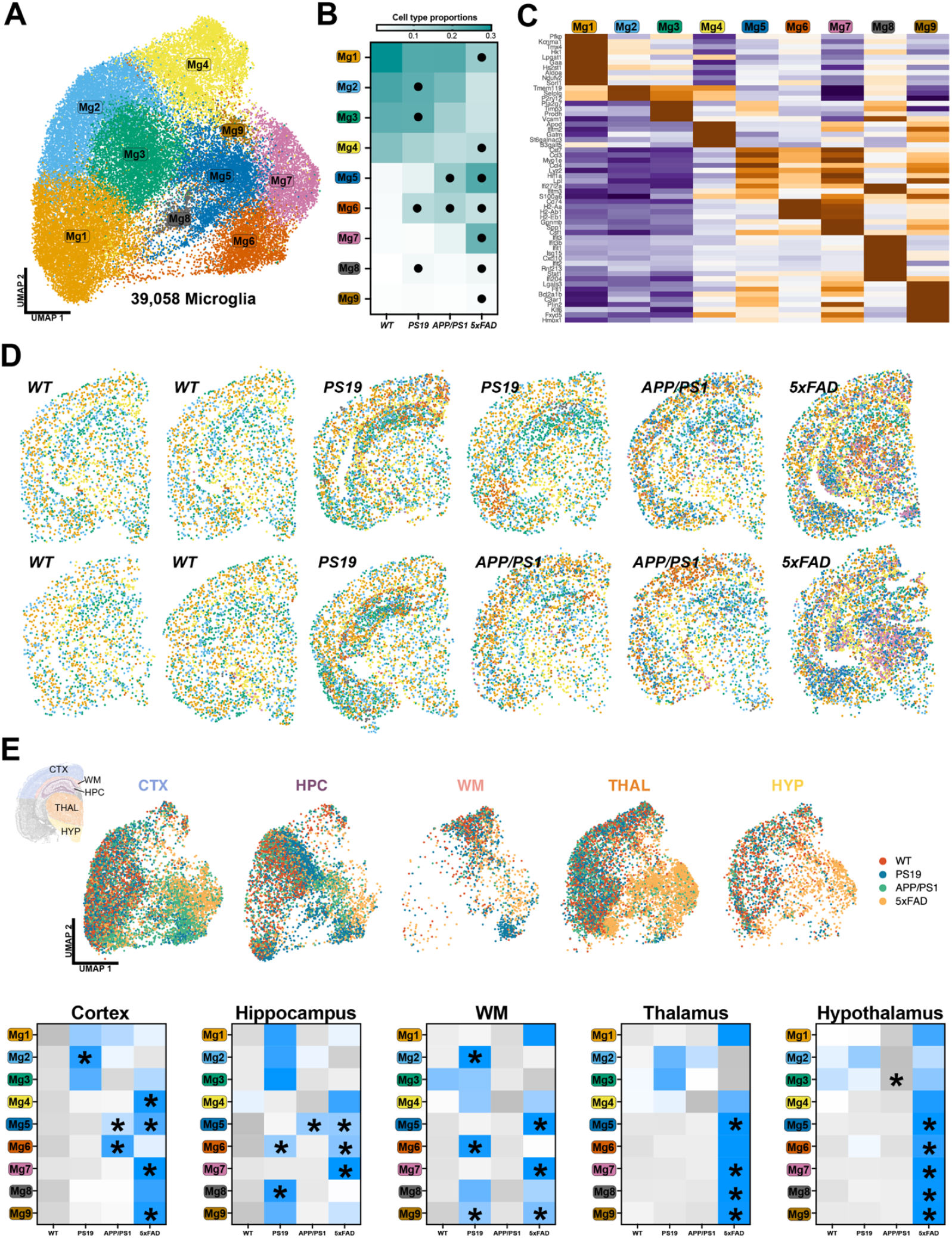
Disease-reactive microglial heterogeneity differentially enriches across anatomical regions and genotype. **A.)** UMAP of 39,058 microglial cells across 9 subtypes from aggregated samples. **B.)** Heatmap on relative cell type proportions per genotype. Black circles indicate *p<0.05 for One-way ANOVA with Dunnet’s posthoc. **C.)** Scaled heatmap of differentially enriched marker genes per microglial subtype. **D.)** Spatial image plots of microglial subtypes across each replicate coronal hemisection. **E.)** UMAP from (A) split by anatomical region of microglia location: cortex (CTX), hippocampus (HPC), corpus callosum/white matter tract (WM), thalamus (THAL), and hypothalamus (HYP), with each colored by genotype (top). Heatmap of z-scores on average counts of each microglia subtype across 5 regions for each genotype (*p<0.05, One-way ANOVA, Dunnet posthoc).

Further, in these five disease reactive states we found similarity in transcriptional profiles with several previously described states from single cell datasets. Mg5 shared close resemblance with “DAM/MgND”-like microglia ^4,5^, Mg6 had the highest expression of MHCII molecules, while Mg7 shared MHCII molecules in tandem with Spp1+/Gpnmb+ closely resembles “ARM”-like microglia ^6^, Mg8 with its high expression of multiple genes linked with interferon responses resembles “IRM”-like microglia ^6,9^, and lastly Mg9 state had enriched expression of several genes related to lipid/cholesterol pathways resembling “LDAM”-like microglia ^8,22^.

After examining transcriptional phenotypes, we subsequently quantified the 9 states across 5 anatomically restricted regions of interest: neocortex (CTX), hippocampus (HPC), corpus callosum/white matter tract (WM), thalamus (THAL), and hypothalamus (HYP). Plotting of the states across the coronal landscapes again demonstrate a stark dichotomy in comparing spatial residence of ‘homeostatic’-like microglia predominating in the WT littermates, while in our 3 AD-like mouse models appearance of Mg5-Mg9 states accumulated in regions continuously implicated in areas of pathological burden (Figure 1D/E). Using the 5 anatomical ROIs, we quantified the average counts of each of the 9 states across all genotypes. In this manner, using the single cell resolving capacity we were able to discriminate the relative abundance of each phenotype. With respect to the disease-reactive states, there were several genotype and region-specific bias in accumulation patterns. PS19 mice displayed varying accumulations across the HPC and WM tract for three states of microglia: Mg6 (MHCII high), Mg8 (IRM-like), and Mg9 (LDAM-like). APP/PS1 mice had significant enrichments across the CTX and HPC of both Mg6 and Mg5 (DAM-like), respectively. As the 5xFAD line has been repeatedly demonstrated to be a very aggressive amyloid plaque depositing model across multiple anatomical areas, our data follow suit for the accumulation of disease-reactive microglial states. Particularly, we demonstrate virtually all microglial states to have significant accumulations across every anatomical region examined, and contrasting to both the PS19 and APP/PS1 models these disease reactive states significantly accumulate in both the THAL and HYP (Figure 1E).

### Proximity to amyloid beta plaques drives differential immunometabolic transcriptional signatures of microglia

As our regional analyses pointed toward accumulation of microglial subsets in anatomically vulnerable areas for AD-related pathology accumulation, we next extended our approach to incorporate distance-based metrics to examine microglial heterogeneity. Using ThioS labeling of amyloid plaques (APP/PS1 and 5xFAD only) in conjunction with co-registration and integration pipelines we dissected the degree that microglial proximity to ThioS-positive amyloid plaques (distance bins of <50 µm, 50-150 µm, and >150 µm) alters transcriptional phenotypes. Our integrative pipeline allowed high precision co-registration of Xenium imaging modalities in conjunction with post-run image capture of ThioS-positive amyloid staining (Figure 3A/B). This procedure then allowed us to apply discrete proximity binning onto each microglial subtype across both APP/PS1 and 5xFAD tissues, we similarly observed a higher proportion of microglia within 50 µm of ThioS+ plaques, compared to APP/PS1 (Figure 3C-E). This spatial characterization recapitulated multiple prior works that have characterized the onset and progression of pathological burden in both cerebral amyloidosis models, wherein the APP/PS1 predominantly acquires amyloid burden in cortical layers and subfield of the hippocampus, compared to significantly more anatomical areas of susceptibility in the 5xFAD ^23^.

**Figure 3.**
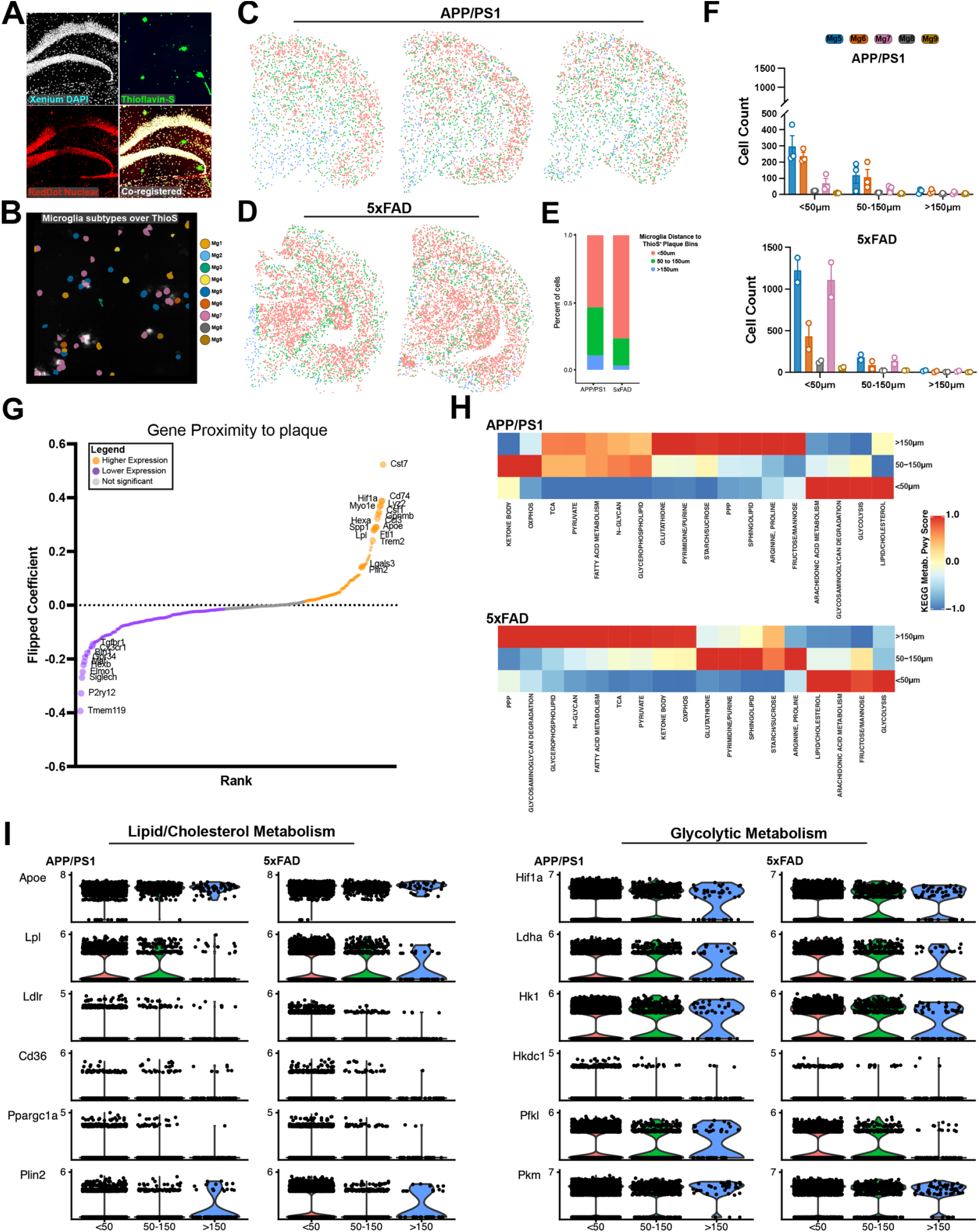
Microglial proximity to ThioS+ amyloid plaques drives altered immunometabolic signatures. **A.)** Representative imaging detailing co-registration of DAPI+ nuclei from Xenium (white) and post-run imaging of thioflavin-s reactive amyloid plaques (green), RedDot nuclei (red) in 5xFAD hippocampus. **B.)** Representative image demonstrating integrated dataset of microglial subtypes overlayed onto ThioS+ plaque image (white). **C/D.)** APP/PS1 and 5xFAD microglia binned upon distance to nearest ThioS+ plaque: red (<50um), green (50-150um) and blue (>150um) across each coronal tissue section. **E.)** Stacked bar plot on distance proportions for APP/PS1 and 5xFAD microglia, demonstrating a higher proportion of microglia in proximity (<50um) compared to APP/PS1. **F.)** Disease-reactive microglia subtype counts separated by ThioS distance bin between genotype, demonstrating a higher proportion of Mg5 and Mg7 subtypes in <50um bin for 5xFAD, relative to APP/PS1. **G.)** Spearman correlation coefficients of gene expression plot as a function of microglia distance to plaque for APP/PS1 and 5xFAD combined, demonstrates higher expression profiles (orange) of multiple disease associated transcripts in tandem with core genes associated with both cholesterol and lipid functions as well as glycolytic response, conversely as distance to plaque increases (purple) multiple genes related to core homeostatic signatures of microglia are increased. **H.)** GSEA pathway analysis of demonstrate proximity-based enrichments of KEGG metabolic pathways across both APP/PS1 and 5xFAD genotypes, wherein similar shifts towards enrichment of cholesterol/lipid and glycolytic pathways are enriched for microglial in closer proximity to ThioS plaques. **I.)** Representative expression plots of genes involved in cholesterol/lipid metabolism and glycolytic metabolism split as a distance function for APP/PS1 and 5xFAD genotypes, collectively demonstrating proximity enrichment.

Comparatively, while both APP/PS1 and 5xFAD had more disease-reactive subsets near amyloid plaques, the distribution of these subtypes within this boundary varied. APP/PS1, while lower in total numbers versus 5xFAD, had relatively equivalent counts of both Mg5 (DAM-like) and Mg6 (MHCII high), while 5xFAD had roughly equivalent distribution of the DAM-like Mg5 and Mg7 (ARM-like) (Figure 3F). We next examined gene expression across all microglial subtypes as a distance function to plaque burden and found that irrespective of genotype or subtype a consistent theme emerged; microglia nearing plaques preferentially upregulated genes involved in plaque-sensing/phagocytosis (*Trem2, Cst7*) and signatures associated with glycolytic (*Hif1a*) and cholesterol/lipid metabolism (*Apoe, Lpl, Plin2*). Conversely, genes previously linked with adult non-disease-exposed signatures (*Cx3cr1, Tmem119, Tgfbr1*) were found with higher expression signatures as distance from plaque increased. Collectively, these proximity-based responses recapitulate several aspects of prior work utilizing population bulk sequencing of amyloid plaque-associated/engulfing microglia ^24,25^. To examine proximity-based signatures in greater detail we utilized GSVA pathway analysis on core KEGG metabolic signaling pathways across all microglia between both APP/PS1 and 5xFAD tissues. Convergent enrichment of pathway signatures between both APP/PS1 and 5xFAD tissues were observed for microglia nearest to plaques for Lipid/Cholesterol, Arachidonic Acid, and Glycolysis (Figure 3H). Similarly, many metabolic pathways were downregulated such as OxPhos, TCA, PPP, Pyruvate, and Fatty Acid, to highlight a few (Figure 3H). We examined representative genes for both lipid/cholesterol and glycolysis pathways and noted a proximity-dose dependency of microglia, such that closer proximity to ThioS-positive plaques was associated with higher gene expression (Figure 3I).

### Human microglia acquire diverse spatial profiles that are linked with proximity to amyloid plaques

To determine the extent that our AD modeling of microglial immunometanolic signatures translates into the ground truth human condition, we spatially profiled 188,624 total cells across dlPFC tissue, encompassing 10 classes of cell types comprised between 12 distinguishable clusters (Figure 4A-E) from a Braak VI autopsy sample. Microglia (TMEM119+) cells were subsequently subsetted and re-clustered, revealing 11,057 total microglia across seven transcriptional states; MG1-MG7 (Figure 5A). Several clusters exhibited expression profiles that mimic disease-reactive phenotypes, particularly MG1, MG3, and MG7. MG1 had several genes that sync with prior work in mouse disease/pathology-reactive phenotypes via expression of *GPBNM, LPL, MERTK*, and APOE. Similarly, MG3 expressed multiple disease/pathology reactive genes but a predominance of MHCII related; *HLA-DQA1, HLA-DRB5, CD74, C1QC, APOE* while MG7 expressed several genes canonically associated with interferon/stress responsive signatures; *IFITM1, IFITM3, STAT1, BCL2L1*. MG2 had expressed the highest levels of genes classically associated with a ‘homeostatic’ phenotype, notably *OLFML3, P2RY12*, and *CX3CR1* (Figure 5B).

**Figure 4.**
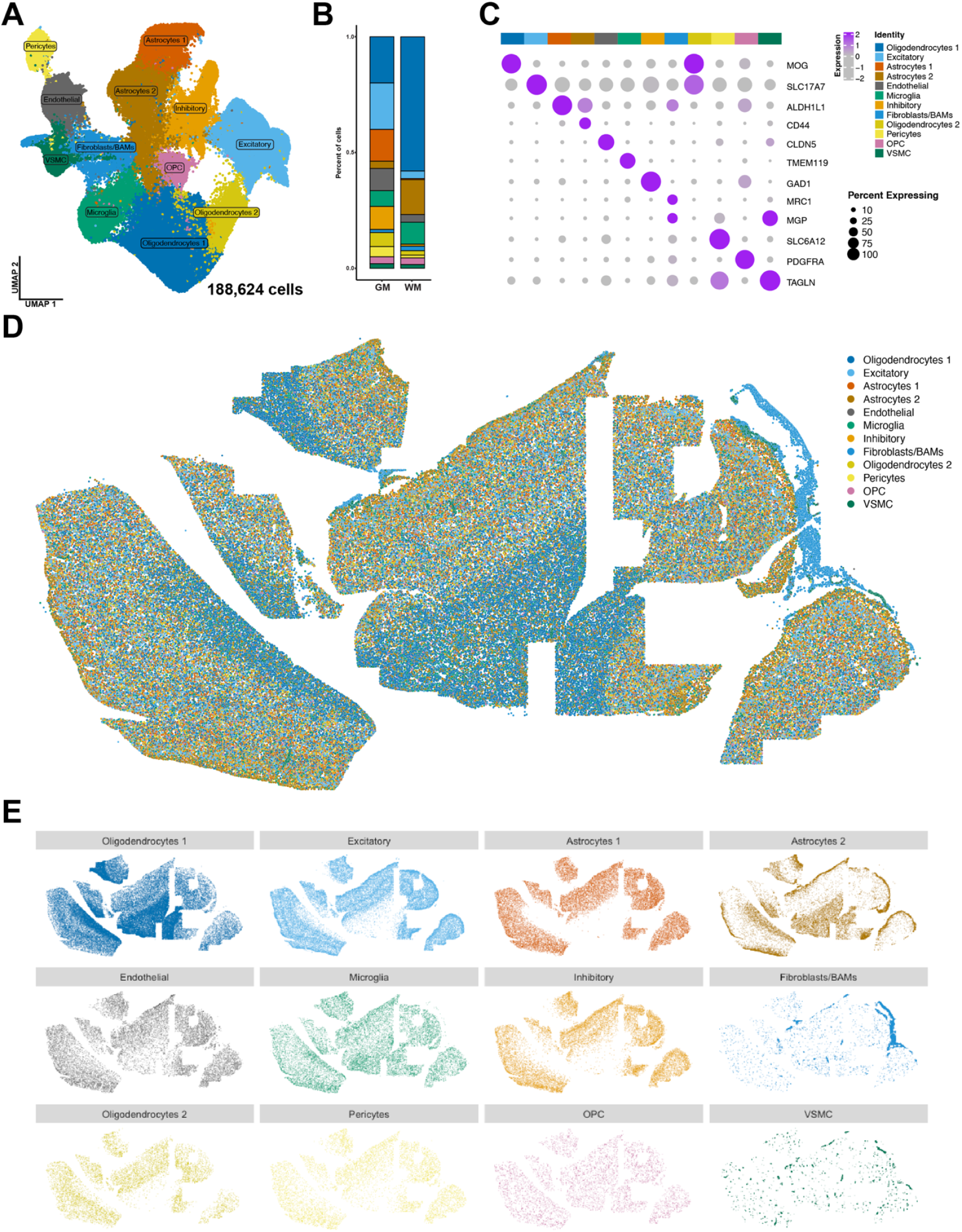
High-resolution mapping of human dlPFC Braak VI tissue. **A.)** UMAP depicting 188,624 total cells across 12 classes of cells in both gray- and white matter **B.)** Stacked bar plot on relative proportions of the 12 cell types split by gray (GM) or white matter (WM) anatomical delineations, with white matter demonstrating a stark enrichment of oligodendrocytes (dark blue), compared to GM. **C.)** Scaled dot plot of canonical cell type marker genes used to define each cluster. **D/E.)** Whole image plot of cell types and split respectively reveals distribution across the entire dlPFC landscape.

**Figure 5.**
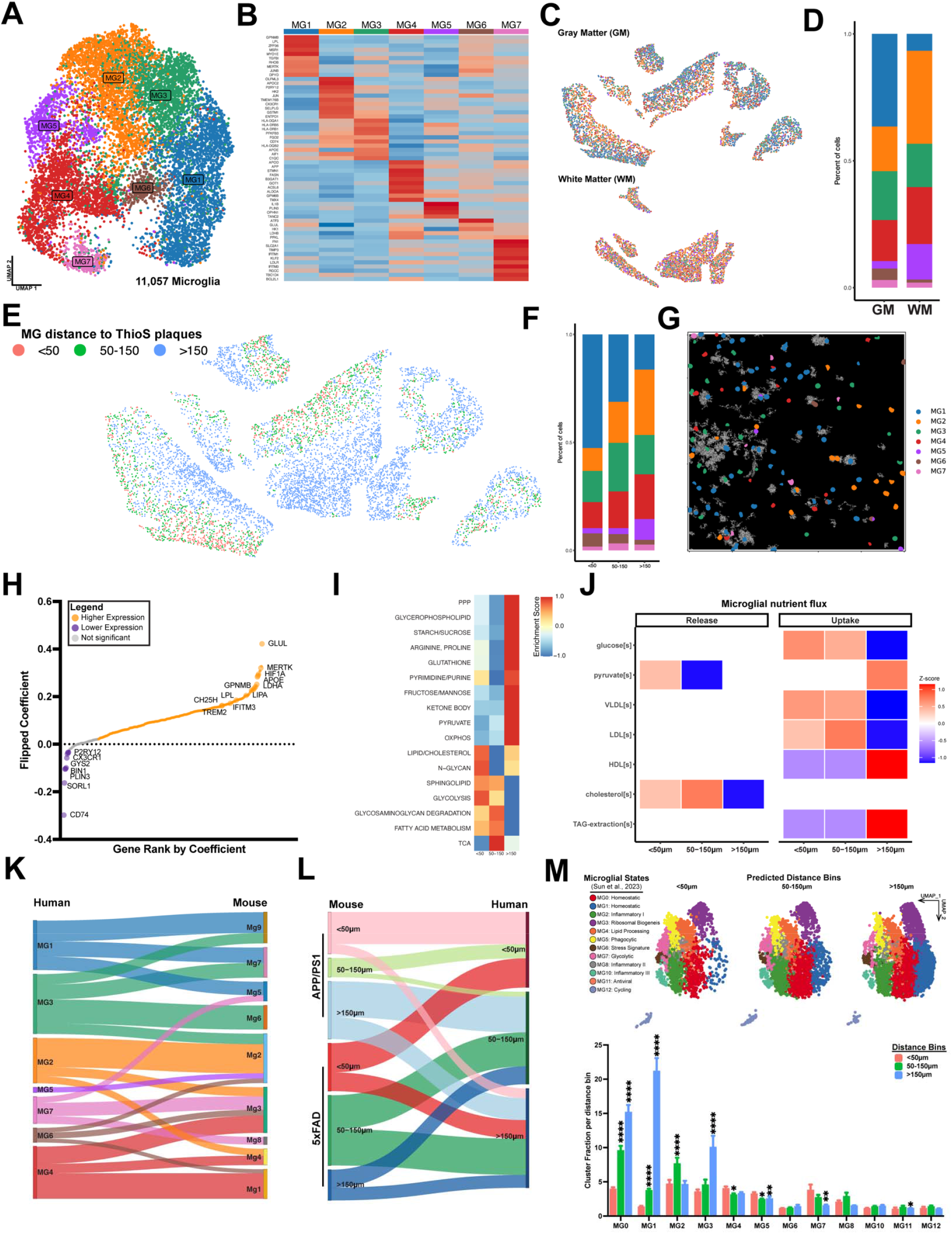
Anatomical location and proximity to amyloid pathology drive human microglial spatial heterogeneity. **A.)** UMAP depicting 11,057 human microglial cells across 7 subtypes from the dlFPC. **B.)** Scaled heatmap of differentially enriched genes per microglial subtype cluster. **C/D.)** Microglial subtypes split by GM or WM anatomical location to which stacked barplot demonstrates the relative proportions across these regions. **E.)** dlPFC microglia as a function of distance proximity to ThioS+ amyloid plaques demonstrates enrichment in GM, while sparse instances of plaque-associated microglia are observed in WM. **F.)** Proportional distribution of MG subtypes as a function of proximity to ThioS+ plaques. **G.)** Representative co-registered ThioS plaque image binarized (white) with MG subtypes overlayed, indicating proximity of MG subtypes, particularly MG1 with amyloid burden. **H.)** Spearman correlation coefficients of gene distance to ThioS+ plaque expression profiles across all MG subtypes, indicating both cholesterol/lipid and glycolytic metabolic profiles that are higher closer to plaques. **I.)** GSVA pathway scoring of selected KEGG metabolic pathways as a function of MG proximity to ThioS plaques. **J.)** METAflux calculation of metabolic reaction activity scores calculated as a distance function to ThioS plaques. Relative nutrient flux categorized between release or uptake indicating MG transcriptional preferences for pyruvate and cholesterol release, contrasting to glucose and lipoprotein particle uptake. **K.)** Sankey diagram of Jaccard similarity scoring for human MG:mouse Mg subtypes demonstrating overlapping features of disease reactive subtypes in human dataset with our mouse subtypes, particularly with MG1and MG3 to Mg5, Mg7, and Mg9. **L.)** Sankey diagram of Jaccard similarity scoring for proximity phenotype of mouse Mg subtypes against human MG subtypes between both APP/PS1 and 5xFAD genotypes. **M.)** Top: Random Forest classifier prediction of cell proximity to amyloid plaque projected onto Sun et al. 2023 PFC microglial nucseq dataset UMAP clusters split by predicted proximity bins. Bottom: Sun et al., MG subtype counts split by projected distance bins reveals shift in “Homeostatic” subtypes (i.e. “MG0”, “MG1”) compared to “Lipid Processing”, “Phagocytic”, and “Glycolytic” (i.e. “MG4”, “MG5”, and “MG7”),”), *p<0.05, **p<0.01,***p<0.001,****p<0.0001 Wilcoxan rank-sum test, Benjamini-Hochberg corrected

We observed that MG1, MG6, and MG7 were found predominantly in the gray matter, whereas MG2 and MG5 were at higher levels in the white matter (Figure 5C/D). Using our proximity to ThioS+ plaques to bin microglia, MG1 and MG6 were found to have the greatest proportional abundance, which proceeded in a stair-step manner of declining abundance as a function of distance (Figure 5E-G). Conversely, MG2, which had the highest expression of genes canonically ascribed as ‘homeostatic’ markers had a comparatively inverse relationship with ThioS and therefore increased in abundance as distance from plaques increased (Figure 5E-G). As in our mouse spatial profiling, we next examined gene expression abundance agnostic of MG clusters and found a similar expression profile dictated by a gene’s proximity to ThioS plaques. Particularly, both disease-reactive markers increased (i.e. *MERTK, GPNMB, TREM2*), while also markers associated with glycolysis (*HIF1A, LDHA*) and cholesterol/lipid signaling (i.e. *LPL, LIPA, CH25H, APOE*) (Figure 5H).

Scoring of all MG clusters/cells against KEGG metabolic pathways, we observed similar features to our mouse modeling work, such that microglia in close proximity (<50um) to ThioS plaques had upregulated signatures for several metabolic pathways, including Glycolysis, Lipid/Cholesterol, Fatty Acid, Sphingolipid. Conversely, these microglia downregulated pathways such as PPP, TCA, OxPhos, Pyruvate (Figure 5I). To gain further insight into the metabolic signatures of our human microglia we employed METAFlux analysis, which generates relative metabolic velocity reactions based upon transcriptional expression profiles. Utilizing our proximity-based bins to ThioS plaques, we demonstrate that microglia have a predicted preferential release pyruvate and cholesterol, while uptake of glucose, and (very)low-density lipoproteins are predicted for nutrient uptake (Figure 5J).

Lastly, we performed several integrative measures to assess the degree of overlapping phenotypes across our mouse dataset as well as prior single-nuclei sequencing from human PFC tissues in the ROSMAP cohort ^26^. Using Jaccard similarly scoring, we examined the degree of overlap between our mouse (Mg) and human (MG) clustering. MG1 showed overlapping features aligning with Mg5 (DAM-like), Mg7 (ARM-like), and Mg9 (LDAM-like). MG3 similarly overlapped with those mouse disease-reactive clusters, while also aligning with Mg6 (MHCII high). Interestingly, MG7 while overlapping with Mg8 (IRM-like) and Mg5 (DAM-like) also intersected with our ‘homeostatic’ Mg3 cluster, likely due to shared *Timp3/TIMP3* expression. MG4, MG5, and MG6 similarly aligned across several of our mouse ‘homeostatic’ clusters (Figure 5K). Next, we assessed human:mouse overlap agnostic of clustering alignment and based upon gene signatures binned by distance to ThioS plaques by genotype (i.e. APP/PS1 and 5xFAD). Predominantly, there was significant signature overlap in our distance bins between both mouse and human across both genotypes (Figure 5L), particularly for <50um in both datasets. Our last method for orthogonal corroboration was to utilize the dataset from Sun et al. ^26^, which characterized microglial phenotypes across six anatomical regions of the human brain, to which we extracted only the PFC nuclei from their dataset. Using a machine-learning approach, we generated predicted distances utilizing our spatial data and projected those onto the signatures/clusters defined by Sun et al. ^26^. These results demonstrated that “homeostatic-MG0, MG1”, and “biogenesis-MG3” all displayed a predicted shift away from pathology (Figure 5M), while “lipid processing – MG4”, “phagocytic – MG5”, and “glycolytic – MG7” were predicted to be higher proportionally nearest to plaques.

### Human disease-reactive microglia associate with select inhibitory neuronal subtypes

Recent work examining microglial reactive morphology in tandem with spatial transcriptomics highlighted a vulnerability of specific inhibitory neuronal populations within the medial temporal gyrus; this vulnerability was dichotomized into ‘early’ and ‘late’ pseudoprogression analyses ^27^. To examine whether there were any correlative relationships with our MG subtypes and neuronal populations we utilized the Allen Atlas in conjunction with deconvolution approach to resolve fine grain mapping of neuronal subtypes. Excitatory and inhibitory populations (Figure 4A) were subsetted and subsequently deconvolved using SpaceXR to reveal 13 neuronal populations spanning both excitatory and inhibitory subtypes (Figure 6A). These subtypes in conjunction with our MG subtypes were analyzed using a predictive neighbor proximity analysis in Squidpy. Our findings demonstrate that both disease-reactive subtypes, MG1 and MG3, were in higher proximity to both VIP+ and Pvalb+ neuronal subtypes, to which MG7 displayed similar proximity, but to a lesser degree (Figure 6B-D). Comparatively, the more ‘homeostatic’ MG2 subtype was positively correlated with several excitatory classes of neurons; Car3+, L6b, L6 CT. This suggests that spatial relationships between distinct microglial states and specific neuronal populations may reflect selective vulnerability or microglial responses tailored to neuronal circuit disruptions in Alzheimer’s disease. Therefore, our data underscore the necessity of spatially resolved approaches to fully understand cellular interactions and circuit-level implications of microglial heterogeneity in neurodegeneration.

**Figure 6.**
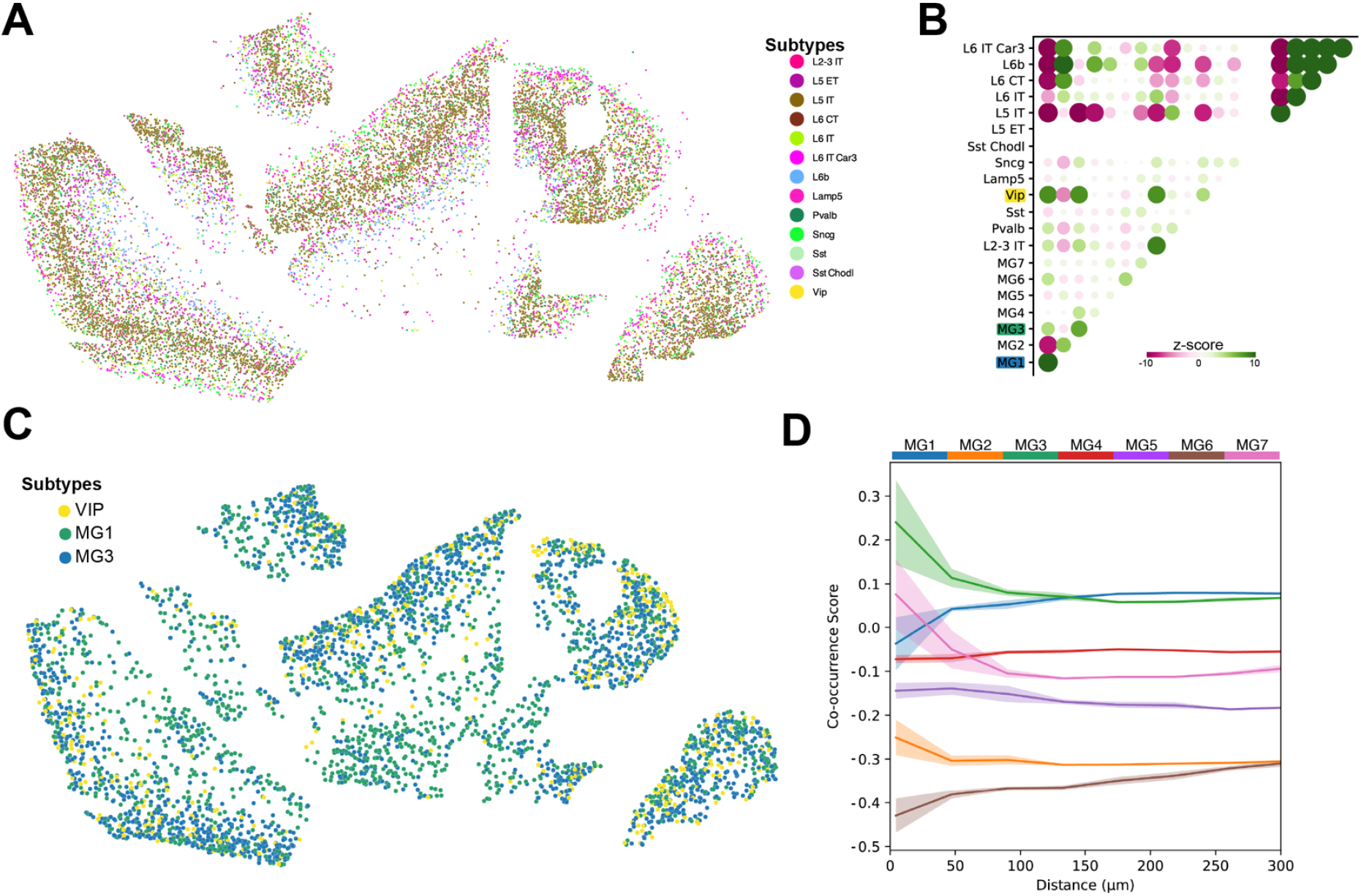
Disease reactive microglial subtypes associate with inhibitory neurons in the dlPFC. **A.)** Deconvolution of neuronal cell types reveal thirteen canonical excitatory and inhibitory subtypes across the dlPFC. **B.)** Spatial neighbor enrichment analysis quantifying whether microglial subtypes are located next to specific neuronal populations more often than expected by random chance. Z-scores derived from permutation testing reveal disease-reactive subtypes MG1 and MG3 preferentially associate with inhibitory neurons including Vip, Pvalb. **C.)** Spatial plot of disease-reactive microglia in tandem with example inhibitory VIP neuronal subtype visualizing proximity relationships. **D.)** Co-occurrence calculations of seven MG subtypes as a function of proximity in microns to VIP neurons, showing the likelihood of finding each microglial subtype near VIP neurons normalized to baseline occurrence. MG1, MG3, and MG7 show elevated co-occurrence scores within the 0-50 micron radius, ± standard deviation.

## Discussion

In the present study, high-resolution spatial transcriptomics was leveraged to investigate microglial immunometabolic adaptations as a function of proximity to AD pathology. We identified nine transcriptionally distinct microglial states (Mg1–Mg9) across mouse AD models (APP/PS1, 5xFAD, and PS19) and seven states (MG1–MG7) in human dorsolateral prefrontal cortex. Microglia proximal to amyloid plaques underwent pronounced glycolytic reprogramming (Hif1a, Ldha) and lipid metabolism shifts (Apoe, Lpl, Plin2), coupled with reduced oxidative phosphorylation and homeostatic marker expression (Tmem119, P2ry12). These are novel findings but also are in accord with rapidly-expanding recent literature related to microglial pathogenetic mechanisms in dementia-related diseases of aging.

Microglia near amyloid plaques robustly upregulated glycolytic genes (Hif1a, Ldha), consistent with metabolic reprogramming described in activated microglia responding to AD pathology ^28,29^. Jung et al. (2025)^30^ emphasized glycolysis as an immediate metabolic response facilitating rapid ATP production under pathological conditions, supported by findings of enhanced glycolytic enzyme expression in APP/PS1 (Guillot-Sestier et al., 2021) and 5xFAD mice ^31^. Similarly, human studies employing positron emission tomography (PET) and single-cell RNA sequencing (scRNA-seq) demonstrated heightened glycolysis correlating with microglial activation around plaques ^32,33^. Our spatial analysis provides exceptional confirmation of these observations at single-cell resolution, highlighting glycolysis as a localized metabolic phenotype within the amyloid plaque microenvironment.

Concurrently, we observed downregulated oxidative phosphorylation and TCA cycle gene signatures, providing strong evidence of a metabolic impairment. Mitochondrial dysfunction and reduced respiratory chain activity have been implicated in AD pathogenesis ^34,35^. Such mitochondrial impairment in microglia contributes to sustained inflammatory states and reduced capacity for Aβ amyloid clearance (Pan et al., 2022; Ulland et al., 2017). Thus, our findings provide spatial resolution to help explain previously reported bioenergetic defects, underscoring glycolytic reliance as both a metabolic adaptation and potential precursor to microglial exhaustion.

Our data revealed marked upregulation of lipid metabolism genes (Apoe, Lpl, Plin2) in microglia surrounding amyloid plaques, consistent with lipid-droplet-accumulating microglia (LDAM) phenotypes described in aging and AD contexts ^8,22^. LDAMs display impaired phagocytosis and heightened inflammatory signaling, exacerbating local neuroinflammation ^36^. Genes such as Apoe and Trem2, strongly associated with lipid handling and plaque interaction ^4,14,37^, were notably elevated near plaques, supporting their role as metabolic orchestrators in reactive microglia.

Prior scRNA-seq studies identified DAM populations enriched in lipid metabolism pathways ^4,5^, while human transcriptomic analyses similarly linked lipid dysregulation to cognitive decline and amyloid burden ^15,26^. Our spatial findings thus directly visualize these transcriptional shifts within specific anatomical niches, providing compelling evidence that lipid accumulation and altered cholesterol dynamics are spatially constrained phenomena driven by proximity to pathological lesions.

Our analysis of human cortex indicated that there are conserved disease-reactive microglial states (MG1, MG3, MG7), paralleling mouse models, reinforcing cross-species validity of AD pathology-associated immunometabolic shifts. Previous human AD studies also report microglial subtypes characterized by enhanced lipid metabolism and inflammatory gene signatures ^14,15,26^. Integration with the ROSMAP human single-nucleus RNA-seq dataset ^26^ further validated transcriptional overlap between spatially-defined and previously described microglial states.

The conservation and specificity of spatially defined microglial metabolic states offer promising therapeutic avenues. Targeting metabolic regulators such as HIF-1α, implicated in glycolytic reprogramming ^28^, or modulating lipid pathways through APOE-TREM2 interactions ^37,38^, presents strategies for restoring microglial homeostasis and mitigating pathological progression. Additionally, leveraging AMPK activators or hexokinase modulators could recalibrate microglial metabolic flux, potentially reversing functional impairment ^29,39^. Importantly, our spatially resolved data indicate therapeutic efforts might be most effective when precisely targeted at metabolic pathways within plaque-proximal microglia rather than generalized immunomodulation.

Beyond the findings of DAM-related gene expression programming in microglia that surrounding amyloid plaques, our approach enabled additional observations related to in situ relationships between cells, gene expression, and pathology. For example, our spatial approach uncovered in situ microscopic intercellular and micro-anatomical vulnerability relationships, including cytological associations between reactive microglia (MG1, MG3) and specific inhibitory neuronal subtypes (VIP+, Pvalb+). Recent evidence highlighted that specific subsets of inhibitory neuronal dysfunctions, particularlya among VIP+ and Pvalb+ interneurons, is integral to early AD pathology ^27^. Thus, our findings suggest that microglial metabolic states may directly influence or reflect neuronal vulnerability, underscoring complex intercellular dynamics within AD.

## Conclusions

Our study spatially contextualizes critical immunometabolic adaptations previously described using bulk and single-cell analyses, elucidating glycolytic and lipid-driven metabolic reprogramming in microglia localized around AD-type pathology (amyloid plaques). By integrating mouse and human datasets, we confirmed the evolutionary conservation and histologic specificity of these metabolic states. This spatial framework enhances our understanding of microglial heterogeneity and highlights metabolic pathways as actionable targets for therapeutic development in AD.

## Methods

### Animals

All experiments were conducted in accordance with the National Institutes of Health Guide for the Care and Use of Laboratory Animals and were approved by the Institutional Animal Care and Use Committee of the University of Kentucky. Three AD-recapitulating mouse models along with their wildtype (WT) littermate controls were employed in this study. Specifically, *P301S* (PS19, 11 months old, n= 3), *APP/PS1* (11 months old, n=3), and *5xFAD* (12 months old, n=2). All mice were group housed 4–5 per cage in individually ventilated cages, in environmentally controlled conditions with standard light cycle (14:10 h light to dark cycle at 21°C) and provided food and water ad libitum.

### Mouse tissue sample preparation for Xenium

Mice were euthanized via rapid cervical dislocation followed by decapitation. Brains were immediately removed, bisected along the midline and further dissected at approximately AP -.80 to -2.50mm from bregma generating a small coronally-oriented tissue chunk. Tissues were subsequently embedded in Tissue-Tek O.C.T. compound (#4583) in a 10×10×5mm cryomold (Tissue-Tek #4565) and submerged in dry-ice chilled (approximately -70°C) isopentane (Sigma #M32631) for exactly 1.5 minutes. Frozen embedded brain tissues were kept on dry ice until being transferred to -80°C storage. Embedded tissues were equilibrated at -20°C for 2hrs prior to sectioning at 10μm using a cryostat (Leica) in the coronal orientation. Coronal hemi sections were placed on Xenium capture slides (10X Genomics, 10006559). Once a capture slide was filled with 6 hemi sections, it was placed in an airtight container and returned to -80°C storage until processed. Slides were processed for Xenium assays using following manufacturer’s suggested protocols (10X Genomics, CG000581RevD, CG000582RevG) in tandem with a custom 433 probe (mouse; Supplementary Table 1). This gene panel consisted of three modules; canonical markers for celltyping, genes associated with microglial/neuroimmune reactivity, and genes associated with cellular metabolism. Immediately following Xenium runs, slides were washed three times with PBS containing 0.05% Tween-20. Subsequently, slides were stained with 0.5% thioflavin-s (ThioS), using standard procedures, before coverslipping with fluorescent mounting media containing NucSpot 640 (Biotium, 23019). Subsequently, slides were imaged using Zeiss AxioscanZ.1 using a 20x objective under standardized acquisition parameter such that all sections were acquired using the same settings. Resultant images were exported using the “ome.tiff” image formatting.

### Human autopsy tissue sample preparation for Xenium

Fresh-frozen dorsolateral prefrontal cortex (dlPFC; Brodmann Area 9) tissue was obtained from the UKY-ADRC autopsy cohort biobank. Details of this community-based cohort, including inclusion/exclusion criteria, neurological assessments, and neuropathological workups, have been published elsewhere ^40,41^. Tissue was obtained from a single female participant who died with documented dementia. Post-mortem interval was 2.9 hours, MMSE = 22, age at death = 68 years, Thal Aβ phase 5, A =3; Braak stage = VI, B=3; and CERAD Frequent, C=3 ^42^. Tissue was embedded in OCT media prior to sectioning on a freezing cryostat. Tissue was sectioned at 10μm and immediately mounted onto the Xenium slide (10X Genomics, 10006559), before storing at -80°C until processed. Slides were processed for Xenium assays using following manufacturer’s suggested protocols (10X Genomics, CG000581RevD, CG000582RevG) in tandem with a custom 479 probe panel (human; Supplementary Table 2). Subsequently, slides were stained with 0.5% thioflavin-s (ThioS), using standard procedures, before coverslipping with fluorescent mounting media containing DAPI (ThermoFisher, P36931). DAPI and ThioS staining on the dlPFC tissue utilizing the “Scan Large Image” function on a Nikon AXR confocal microscope using a 20x objective. Resultant images were exported using the “ome.tiff” image formatting.

### Xenium assay: data processing

Xenium raw data were processed using XeniumRanger (v2.0.1.2; mouse) and XeniumRanger (v3.0; human), with segmentation set to 5μm. Sections were annotated by genotype and replicate via XeniumExplorer (v2.0) ‘freehand’ selection tool and the resultant CellIDs were exported. Seurat (v5.1) was used for initial object creation, removing cells with “0” counts ^43^, and merging of the two slides (mouse) into a single Seurat object were completed. PCA and UMAP were used for two-dimensional visualization of the dimensionally reduced dataset (with the top 30 PCs used, based upon the total variance explained by each), *FindClusters* with Louvain algorithm set at 0.3 resolution. Subsequently, *FindMarkers* function was executed with Wilcox algorithm to generate Top10 cluster differentially expressed genes (DEGs) to enable celltyping. This process yielded 797,255 cells across 16 clusters for mouse and 188,624 cells across 12 clusters for human respectively. Canonically defined CNS cell type markers were used to identify cell types in the mouse model which include: *Tmem119* (Microglia), *Snap25, Gad1, Pvalb, Sst* (Inhibitory neurons), *Cd3e* (Lymphocytes), *Emcn* (Endothelia), *Aldoc* (Astrocytes), *Snap25, Slc17a7, Lamp5* (Excitatory neurons), *Pdgfra* (Oligodendrocyte precursors), *Mog* (Oligodendrocytes), *Kcnj8* (Pericytes), *Bgn* (Vascular leptomeningeal), *Car12* (Choroid plexus), and *Tmem212* (Ependymal). Simultaneously, CNS cell type identification was performed on the human model using canonical markers that are orthologous to those employed in the mouse model, such as: *MOG* (Oligodendrocytes), *SLC17A7* (Excitatory neurons), *ALDH1L1* (astrocytes), *CLDN5* (endothelia), *GAD1* (inhibitory neurons), *CD44 and MRC1*(Fibroblasts/Border associated macrophages), *SLC6A12* (Pericytes), *PDGFRA* (Oligodendrocyte precursors), *TAGLN* (Vascular smooth muscle). Following CNS cell typing, microglial populations for both mouse and human tissues were subsetted into new objects retaining only immunometabolic transcripts, resulting in 339 and 384 total genes in each set, respectively.

Following identical processing as used above via SCTransform, PCA, UMAP, and FindClusters (0.3 resolution for mouse and 0.35 resolution for human) yielded 39,050 microglia across 9 clusters for the mouse model and 11,057 microglia across 7 clusters for the human model. Granular neuronal subtypes were further identified using spatial deconvolution in SpaceXR (v2.2.1) with the Allen cortical brain reference (https://portal.brain-map.org/atlases-and-data/rnaseq/human-m1-10x) in doublet mode.

### Proportional calculations

For the mouse model, proportions of each cell type were calculated between each genotype and then graphically represented using a heatmap. Raw counts of each cell type were obtained per genotype, and subsequent one-way ANOVA (base R) and dunnet posthoc analysis (R package multcomp version 1.4) were used to determine statistical significance among cell type distribution between genotypes. Identical statistical measures to determine microglial subtype distributions across mouse brain regions.

### Thioflavin-S image co-registration and data integration into Xenium objects

Scanned ome.tiff images containing ThioS were co-registered to Xenium DAPI image via QuPath (v0.5.1) utilizing Fiji (ImageJ 1.54f) with BigDataViewer-Playground BDV Dataset. Warpy in Fiji with DAPI (Xenium image) was set as the fixed source and NucSpot 640 (mouse) or DAPI (human) was set as the moving source. Registration was achieved using BigWarp with 30 landmarks per tissue section for the mouse model and 160 landmarks total for the human tissue. The resultant co-registered image was exported with Warpy transform and imported into QuPath via Warpy Image Combiner. Python skimage (0.22.0) package was utilized to split channels of the co-registered image to allow a single channel (e.g. ThioS) to export into a format that was compatible with HALO image analysis software (v4.0). Within HALO, images were annotated again by genotype and replicate for all APP/PS1 and 5xFAD tissues, or as ‘white’ or ‘gray’ matter ROIs for human tissue. ThioS positive pixels were defined using *Multiplex Co-localization* algorithm in HALO, with ThioS objects’ location stored as XY coordinates. ThioS coordinates were imported into a pandas dataframe, where centroid calculations and pixel-to-micrometer conversions were performed. Using Thios centroid of each plaque, the minimum Euclidean distance between each cell and plaque was calculated using distance.cdist in scipy Python software package (v1.15.1). The minimum distance was then stored as metadata, re-imported into the Seurat object, and categorized into <50µm, 50-150µm, and >150µm bins for downstream analysis.

### Pathway scoring

To identify gene signatures at the single-cell level, we used the Escape R package (v2.2.3) with the GSVA method (v2.1.5) under default normalization settings. Pathway scoring was performed using a curated list of genes of rate limiting and canonical markers associated with gene sets for selected KEGG metabolic pathways. Scores were calculated for individual cells, and their distribution across distance bins was analyzed to assess gene signature expression as a function of proximity to plaques.

## Visualization

Python software package spatialdata (v.0.3.0) was used to superimpose images of microglia over ThioS. To accurately superimpose the images, coregistered ThioS images from above were imported, and initially adjusted for contrast and brightness. Subsequently, an added channel dimension was then introduced to each image using numpy.expand_dims(), parsed using Image2DModel.parse, and then validated using Image2DModel.validate() from spatialdata. The images were then added to spatialdata objects along with the microglial subset information by importing in R Seurat metadata. The final images were cropped using spatialdata.bounding_box_query.

## Gene to distance to plaque characterization

To characterize the cell-type agnostic relationship between gene expression and the distance to plaque, Spearman’s correlation coefficients were calculated for each gene′s expression values against the minimum distance to plaque using the spearmanr function from the SciPy package (v1.15.1) in Python. All genes were subsequently ranked by inverse coefficient to elicit a positive value to reflect higher gene expression as a function of plaque distance. The top 3 positive and top 3 negative values were labeled. Additionally, canonical disease associated microglial genes and hallmark metabolic genes were labeled for further characterization. Python package Matplotlib (v3.10.0) was used for visualization of the generated data.

### Metabolic flux calculation

Relative metabolic flux scores were calculated using the genome-scale metabolic model in METAflux (v0.0.0.9). Metabolic reaction activity scores were derived from gene expression data, and flux calculations were performed across distance bins using human blood as the medium. Flux scores were classified as either nutrient uptake or release depending on their sign. The absolute values of the flux scores were converted to Z-scores and visualized using a heatmap.

### Mouse to Human comparisons

To determine similarity between mouse and human clusters, DEGs were calculated using Seurat FindAllMarkers Wilcox with min.pct of 0.25 and logfc.threshold of 0.25. The 50 most differentially expressed genes per group were selected and compared using GeneOverlap 1.42.0 to calculate Jaccard similarity scores. R software package networkD3 (v.3 0.4) was used to visualize the top two scores for each cluster as a Sankey diagram.

### Predicting distance to plaque on outside datasets

To demonstrate that gene expression levels can be used to infer distance to plaque, we developed a random forest classifier utilizing the Python package scikit-learn (v1.6.1). To address batch effects between the two datasets, we employed the combat function within scanpy (v1.10.4). The model was built using only matching features (313 genes). We split the microglial xenium spatial data into training (75%) and testing (25%) sets, achieving an accuracy rate of 53% on the test set. The trained model was then applied to predict the distance to plaque bins in the microglial data from Sun et al., and the results were projected onto their original UMAP. Only the PFC and ‘late AD’ nuclei were utilized.

### Proximity Analysis

Neuronal and microglial subtypes then underwent proximity analysis using Python package squidpy (v.1.6.2) spatial_neighbors, then nhood_enrichment with generic coord_type. The nhood_enrichment function calculates an enrichment score by evaluating the proximity of cell clusters within the connectivity graph. The observed number of events is compared to a distribution obtained from randomized permutations, and a z-score is computed to assess the statistical significance of the enrichment.

## Acknowledgements

This work was supported by the National Institute of Health, NIA and NINDS (SLM – R01AG068330; PTN - P30 AG072946, P01 AG078116, R01 NS118584, LAJ - R01AG062550, R01AG081421, R01AG080589; and JMM - R01AG070830 and RF1NS118558), the CNS Metabolism COBRE P20 GM148326 (JMM, SLM, and LAJ), BrightFocus Foundation (SLM – A20201775S) and the Alzheimer’s Association (LAJ, JMM).

